# Transport Cocktails for Cancer Therapeutics

**DOI:** 10.1101/2024.01.23.576806

**Authors:** Michail E. Kavousanakis, Omkar Bhatavdekar, Remco Bastiaannet, Yannis Kevrekidis, Stavroula Sofou

## Abstract

Beyond biological cell heterogeneity, evidenced by different resistances to therapeutics, “delivery heterogeneity” crucially limits treatment efficacy for advanced solid tumors: variations in therapeutic drug delivery to different tumor areas (perivascular, perinecrotic) leading to nonuniform drug concentrations/doses and to unsuccessful treatment (cancer cell kill). Short-range (40-80 µm), high energy (1-5 MeV) alpha-particles successfully address the biological heterogeneity: the double-strand DNA breaks they cause make them impervious to cell resistance mechanisms. Multiresponsive nanocarriers and/or engineered antibody-drug-conjugates are elegant approaches to delivering such alpha-particle emitters. Delivery heterogeneity, however, remains a challenge in established (i.e. large, vascularized) tumors. Remarkably, delivery properties enabling efficacy at the cell scale (targeting selectivity, affinity, cell drug uptake) may act against spatial delivery uniformity at the tumor scale (binding-site barrier effect^1^). We have previously demonstrated, in different mouse models, that spatial delivery uniformity, key to the effective killing of solid tumors, can be achieved utilizing combinations of different, distinct delivery carriers of the same emitter, but with different, complementary delivery properties, “leaving no cancer cell behind”. We build first principles reaction-transport models (quantitatively informed by experiments) that explain the “geographically complementary” behaviors of such carrier cocktails, and help optimally design these cocktails and their delivery protocols.

## Introduction

Although cancer death rates are decreasing, mainly due to early detection, cases of advanced metastatic and/or recurrent solid cancers still have no cure^2^. At the advanced stage of established (i.e. large, vascularized) tumors, current therapeutic approaches necessarily resort to combinations of therapeutics^3^; unfortunately, even with highly toxic combination regimens, the vast majority of patients fail to reach a durable response. A major reason for this outcome is the heterogeneity in drug delivery within established solid tumors^4–6^. Most therapeutic agents require being physically present in the vicinity of their molecular target in order to act. The limited penetration into solid tumors of nanocarriers and/or of targeting antibody-drug-conjugates, inevitably leave tumor regions exposed to too low/non-lethal levels of therapeutics, ultimately enabling disease recurrence^4,7–9^. A treatment strategy addressing delivery heterogeneities in established lesions is critical to successfully treating solid tumor patients especially at advanced stages of their disease.

This work is motivated by the *in vivo* observations in Fig. 1^5,10^. The 3D diagram quantifies how, and the representative alpha-camera snapshots above it partially explain why, delivery of *the same injected radioactivity* to vascularized solid tumors in mice *is markedly more successful* in inhibiting tumor growth *when split in equal parts between two separate and distinct carriers*: ∼50% via targeting antibodies and ∼50% via tumor-responsive liposomes. Compared to the same total injected radioactivity delivered either (a) 100% via antibodies, or (b) 100% via liposomes, the “carrier cocktail” performs quantifiably better. The images in Fig. 1 strongly suggest that *this improvement is due to the better tissue penetration of alpha-particles delivered via the carrier cocktail*, in stark contrast to the nonuniformity observed with either pure carrier.

**Fig. 1:**
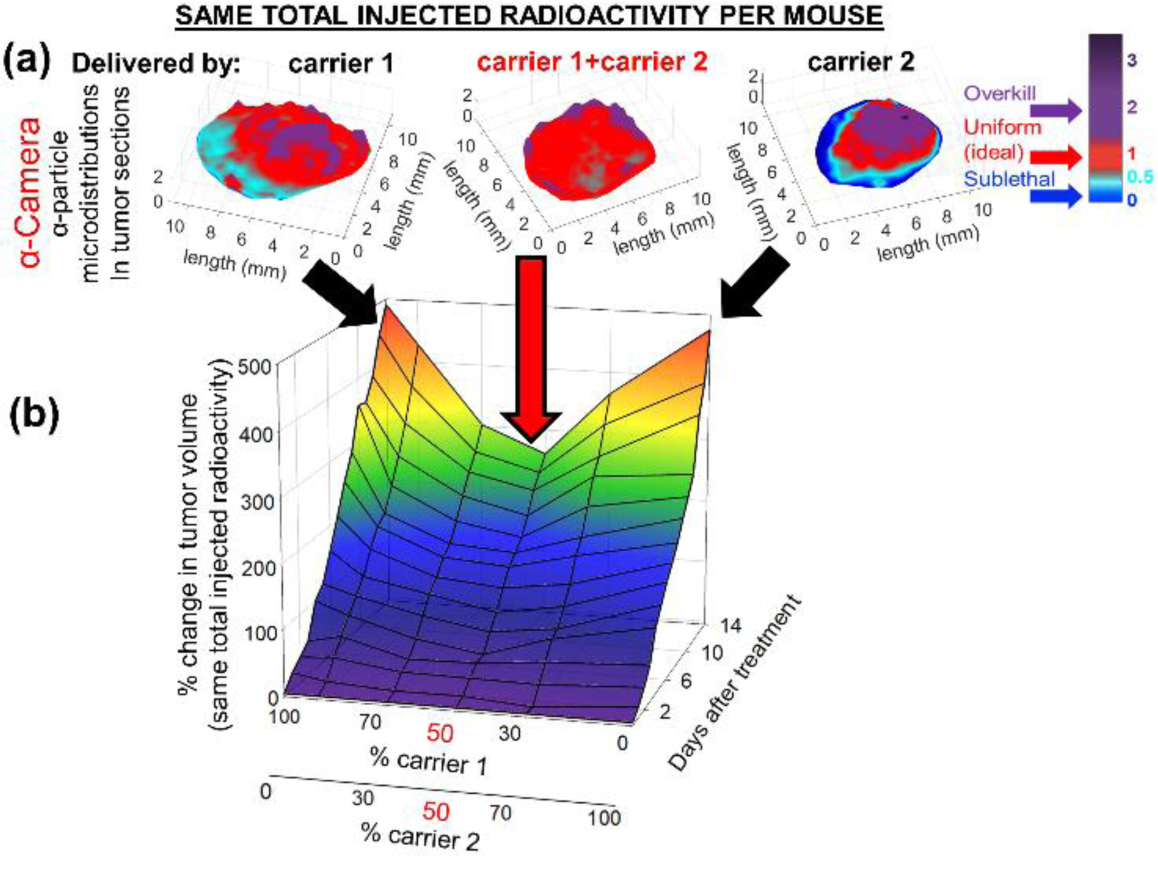
Improved drug spreading within solid tumors using simultaneously two separate delivery carriers.

We set out to quantitatively model these non-intuitive results through first-principles modelling of the transport and kinetic processes involved in the drug delivery process; the kinetic and transport parameters in the mathematical model were arrived at via targeted experiments.

Our two separate carriers have complementary delivery properties: one acting at the cell scale (antibody targeting) and the other acting at the tumor scale (liposome content release). Each carrier type preferentially kills a different region of the tumor ^5,10^: (1) the tumor-responsive liposomes, that, upon tumor uptake, release in the interstitium a highly-diffusing form of their payload, which then penetrates the deeper parts of tumors where antibodies do not reach; and (2) the separately administered, less-penetrating drug/isotope-labeled targeting antibody, that effectively kills the tumor perivascular regions from where the liposomes’ released contents clear too fast.

To improve drug spreading within solid tumors, two separate delivery carriers *of the same drug*, were employed. The carriers were chosen to deliver their therapeutic cargo in complementary regions of the same solid tumor. Collectively, the drug became more well-spread within established, soft-tissue solid tumors, resulting in better tumor growth inhibition. **(a)** Top panel: Spatial microlandscapes (color maps) exhibiting the deviation from uniform irradiation of tumor sections. α-camera images of tumor sections, where pixel intensities of delivered radioactivity were divided by the mean intensity, which was averaged over the entire tumor section, with the aim to reveal deviation from the mean (colored vertical bar). On tumor sections, local α-particle activities that were similar to the mean tumor uptake were colored in red^5^. **(b)** 3D-plot: Tumor growth inhibition was greater when the same total administered activity was equally split between the two separate carriers^10^, even though the tumor absorbed dose at 50:50 split (0.23±0.02 Gy) was lower than the dose (0.34±0.04 Gy) delivered by the radiolabeled targeting antibody alone (100% carrier 2).

Our tumor-responsive liposomes are composed of membranes forming phase-separated lipid domains (resembling lipid patches) with lowering pH^1,11^. During circulation in the blood, such liposomes comprise well-mixed, uniform membranes that stably retain their encapsulated contents. In the acidic tumor interstitium (pHe∼6.7-6.5) lipid-phase separation results in formation of *lipid patches (rafts)* that span the bilayer, creating transient lipid-packing defects along the patch boundaries, and enabling release of encapsulated agents. The liposomes may also exhibit an adhesive property that enables them to bind to the tumors’ extracellular matrix, delaying their clearance from tumors^7,9^.

The antibody-isotope-conjugates utilized herein comprise FDA-approved antibodies, which are designed to stably retain their therapeutic cargo^5,10,12^. These conjugates exhibit strong binding to cell surface markers and become internalized. Currently, alpha-particle radiolabeled antibody conjugates are evaluated in clinical trials against solid tumors of variable origin and over a range of targeted receptor expression. This suggests that our cocktail approach may be broadly applicable.

In this work, we develop and implement an experimentally informed mathematical model that can ultimately predict the best possible combinations of the two carriers for given tumor sizes. Intratumoral spatio-temporal profiles of the therapeutic agents, delivered by each carrier type, and the corresponding intratumoral delivered radioactivity are calculated, so as to identify the optimal delivery modality combination including the associated temporal dosing scheduling.

## Results

We develop experimentally informed reaction-diffusion models to describe isotope delivery jointly from both isotope-releasing liposomes (equations (2)-(5)) and antibodies (equations (6)-(9)). Each model component succinctly captures the distinct transport characteristics of its respective carrier:

a. Tumor-responsive (due to tumor acidity) liposomes release their payload, which can reach the deeper parts of tumors;
b. Isotope-labeled antibodies, though less penetrating, exhibit highly effective tumor cell elimination, particulary in the perivascular areas.

Finally, we systematically explore the optimal synergy between of these carriers, tailoring them to specific tumor sizes.

### Modelling tumor-responsive liposome isotope delivery

For our simulations, we assume a specific activity 0.34 *MBq*/*μmol* lipid^10^. Considering the half-life of ^225^*Ac*, 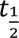 = 9.9 days, and that each decay of ^225^*AC* generates four alpha particles and three radio-active daughters^13^, the radioactivity of ^225^*Ac* is ∼2,000 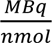 of isotope. Thus, the ratio of isotope content per liposome (considering 100,000 lipids per liposome) is equal to *N_c_* ≈0.017 mol ^225^*Ac* atom per mol liposome. In addition, we assume an apparent diffusion coefficient for the released isotope equal to, *D_C_* = 800 *mm*^2^ · *s*^−1 14^. To obtain the spheroid volume fraction that is accessible to isotope, *φ_c_*, we estimate the volume of the spheroid unoccupied by cancer cells. With a mean diameter of 10 μm per cancer cell, in a spheroid of radius ∼200 μm, that consists of ∼16400 cancer cells, we estimate *φ_c_* ≈ 0.73.

We simulate the incubation of tumor spheroids in a solution of liposome concentration, [*L^(sol)^*] = 30 *μM* for 6 hrs and compute the isotope concentration during isotope uptake, and then during the first 4 hrs upon the completion of incubation (clearance experiments). The parameter values for the liposome-carriers simulation are as follows: the effective diffusion coefficient of liposomes in the spheroid, *D_L_* = 1.5 × 10^−13^ *m*^2^ · *s*^−1^; the mass transfer coefficient for liposomes during uptake and clearance experiments, respectively: *P_L,up_* = 1.9 × 10^−9^ *m* · *s*^−1^, *P_L,cl_* = 5.8 × 10^−9^ *m* · *s*^−1^. These parameter values, used in the simulations, are derived by fitting the model to targeted experiments described in the Methods section. The same set of parameter values is then used for the cocktail simulations.

Liposomes release their isotope content at a rate dependent on *pH*; in particular, the release rate constant, *k_r_*, has been experimentally measured to correlate with *pH* in a linear fashion, *k_r_* = *a* + *b* · *p*. For the purpose of our simulations, we compute the values of constants, *a* and *b* by fitting the isotope release kinetics of liposomes when loaded with a drug-surrogate (see Supplementary Information, Sec. B), and obtain the following simple empirical relation:

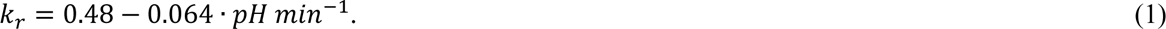

Importantly, experimental measurements show a spatial variation of *pH* within a growing spheroid. Fig.(S-1) of the Supplementary Material depicts the *pH* variation in BT-474 spheroids with the distance from the spheroid’s center. The environment in the tumor’s interior is more acidic, implying higher drug-release rates (see equation (1)), compared to the exterior regions of the spheroid.

Figure 2(A) shows different temporal snapshots of the spatial distribution of liposome concentration within a spheroid following incubation. Liposomes diffuse and manage to penetrate the spheroid up to certain depths during the incubation stage. Even upon completion of isotope uptake, liposomes can still be found at the interior of the spheroid; these are mostly liposomes that have already released all their isotope-content (see Fig. 2(C)). Liposomes will gradually release their isotope as they diffuse towards the interior of the spheroid (and the environment becomes more acidic); one can thus observe higher concentration of lower-content liposomes closer to the tumor’s center. We compute the distribution of isotope, carried and released by the liposomes, during incubation/uptake as well as during clearance phases with respect to the spheroids (Fig. 2(B)). The isotope is released by the liposomes and diffuses quickly towards the interior region of the spheroid, where it remains even after the completion of incubation. Figure 2(D) shows the uniform distribution of the released -by the liposomes-isotope (in its free form). Higher concentration is observed at the outer region of the spheroid, where liposomes with higher isotope content can be found.

**Fig. 2:**
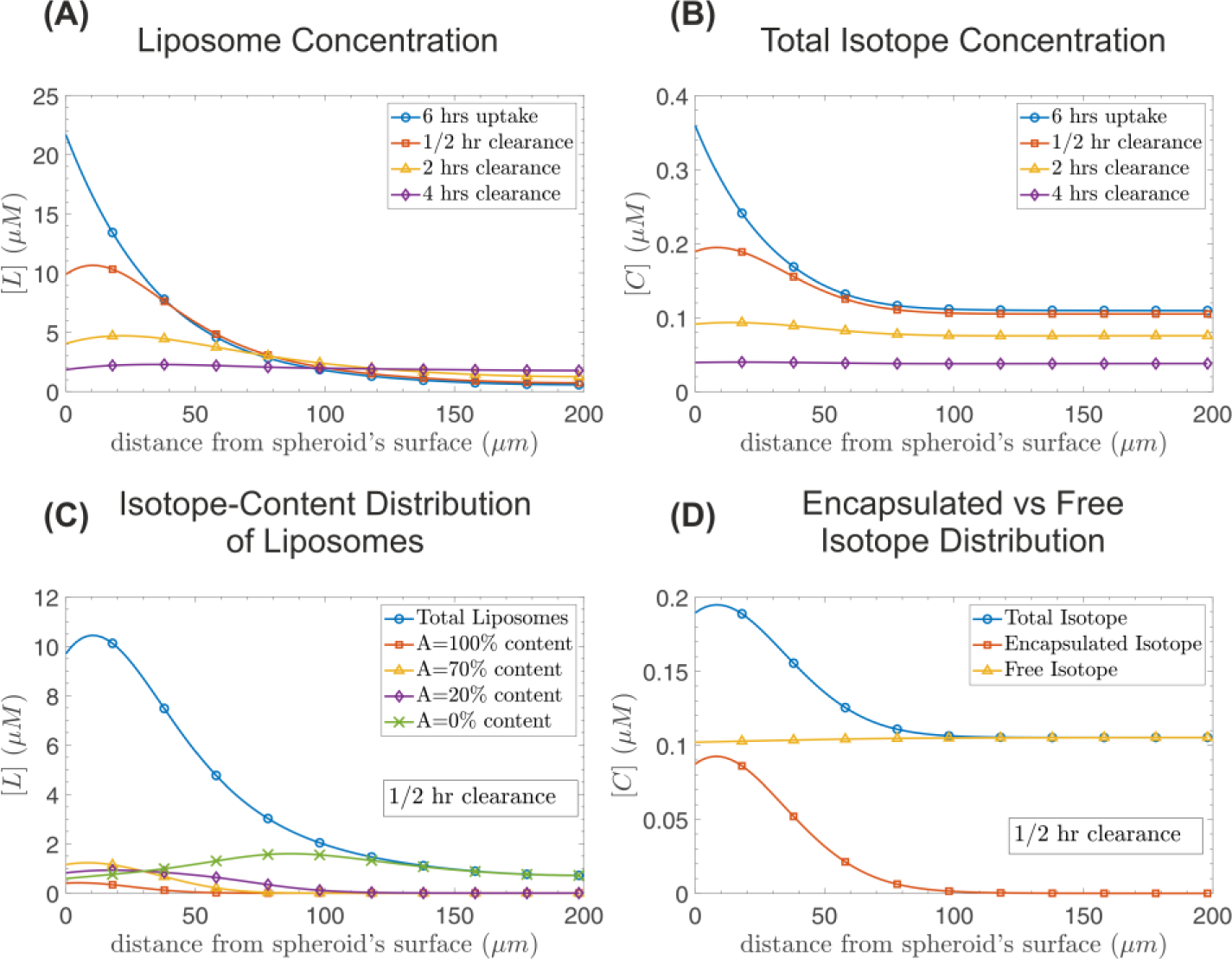
Representative liposome isotope carrier simulation results. (A) Spatial distribution of liposomes in a tumor spheroid at the end of incubation (6 hrs: blue line with open circles), 1/2 hr after incubation (red line with open squares), 2 hrs after incubation (yellow line with open triangles), and 4 hrs after incubation (purple line with open diamonds). **(B)** Isotope concentration spatial distribution at the end of incubation (blue line with open circles), and 1/2, 2, 4 hrs after the completion of incubation depicted with red line and open squares, yellow line and open triangles, and purple line with open diamonds, respectively. **(C)** Distribution of liposomes depending on their isotope content, *A*, 1/2 hr after the completion of the 6-hr incubation experiment. The total concentration of liposomes is depicted with a blue line with open circles; liposomes with *A*=100, 70, 20 % isotope content are shown with red line and open squares, yellow line and open triangles, and purple line with open diamonds, respectively. Liposomes already empty of isotope (∼0 % content) are denoted with the crossed green line. **(D)** The total isotope concentration spatial distribution 1/2 hr after the completion of incubation is shown as a blue line and open circles. The encapsulated isotope in the liposomes is depicted with the red line and open rectangles, while isotope released in the interstitium is illustrated with the yellow line and open triangles. Tumor spheroids are incubated for 6hrs in a solution of liposome concentration, [*L*^(*sol*)^] = 30 *μM*. Parameter values for the simulation: the effective diffusion coefficient of liposomes in the spheroid, *D_L_* = 1.5 × 10^−13^ *m*^2^ · *s*^−1^; the mass transfer coefficient for liposomes during uptake and clearance, respectively: *P_L,up_* = 1.9 × 10^−9^ *m* · *s*^−1^, *P_L,cl_* = 5.8 × 10^−9^ *m* · *s*^−1^; the apparent diffusion coefficient of the isotope, *D_C_* = 2 × 10^−11^ *m*^2^ · *s*^−1^, and the isotope’s mass transfer coefficients, *P_c,up_* = 1.7 × 10^−8^ *m* · *s*^−1^ and *P_c,ci_* = 2.9 × 10^−8^ *m* · *s*^−1^, respectively.

### Modelling antibody isotope delivery

Tumor spheroids are incubated in a solution of constant antibody for 24 hrs *in* silico, as in experimental practice. Antibodies then penetrate the spheroid, bind on the cancer cells, and gradually become internalized by them. We simulate these processes by numerically solving model equations (6)-(9) below. We consider a constant concentration of antibodies in the solution where spheroids are immersed; during the uptake experiments, which last 24 hrs, the concentration of antibodies in the incubating solution is fixed to [*Ab*^(*sol*)^] = 240 nM. If we assume a specific activity 1.87 *MBq*/*mg* Ab^10^ (here the antibody is Trastuzumab with molecular weight 150 *kDa*), the ratio of isotope per Ab is approximately 1.45 × 10^−4^ mol isotope / mol Ab (as reported above the radioactivity for ^225^*Ac* is approximately 2,000 *MBq*/*nmol* isotope). For our simulations, we adopt the following set of parameter values: the effective diffusion coefficient of antibodies in the spheroid: *D_Ab_* = 6 × 10^−12^ *m*^2^ · *s*^−1^; the equilibrium dissociation constant for antibodies: 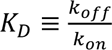 = 5 nM; the dissociation rate constant: *k_off_* = 4 × 10^−3^ *s*^−1^; the internalization rate constant: *k_int_* = 1.4 × 10^−5^ *s*^−1^; the mass transfer coefficient for antibodies during uptake and clearance experiments, respectively: *P_Ab,up_*, = 2.5 × 10^−10^ *m* · *s*^−1^, *P_Ab,cl_* = 8 × 10^−7^ *m* · *s*^−1^; the concentration of unoccupied by antibodies receptors: *R_T_* = 1060 nM. These values are derived from targeted experiments described in the Methods section.

Antibodies bind on the surface of cancer cells, preventing their further transport towards the interior of the spheroid (the binding-site barrier effect ^15–17^). Our model predicts the spatiotemporal profiles of antibodies found in the interstitium ([*Ab*^(*I*)^]), the antibody/receptor complex ([*Ab*^(*b*)^]), and the antibody concentration ([*Ab*^(*int*)^]) internalized in cancer cells. Thus, one can infer the isotope concentration, given that the isotope/antibody molar ratio, as derived from the measured specific activity^5,10^, is 1.45 × 10^−4^. Snapshots of the total isotope concentration (denoted with *C*) are presented in Fig. 3(B). One can observe that the total isotope concentration does not fall below 50% during the first 4 hrs of the clearance experiments; in contrast, when carried by liposomes, we observe a decay at a much higher rate (see Fig. 2(B)). This behaviour can be attributed to the antibodies binding on the cancer cell surfaces, and subsequently becoming internalized. In particular, by breaking the total isotope concentration down to its various constituent forms, we can clearly see that the isotope can mainly be found in the form of antibody/receptor complexes (*C*^(*b*)^), and in internalized antibodies (*C*^(*int*)^). The concentration of antibodies, and thus their isotope cargo, when they diffuse in the spheroid’s interstitium, is at a substantially lower level compared to the other forms of antibodies, during both the uptake and the clearance experiments from spheroids (Fig. 3(C) & (D)). Finally, we observe that the isotope concentration is practically negligible at the inner regions of the spheroid (practically negligible at distances up to 75 μm from the spheroid’s center).

**Fig. 3:**
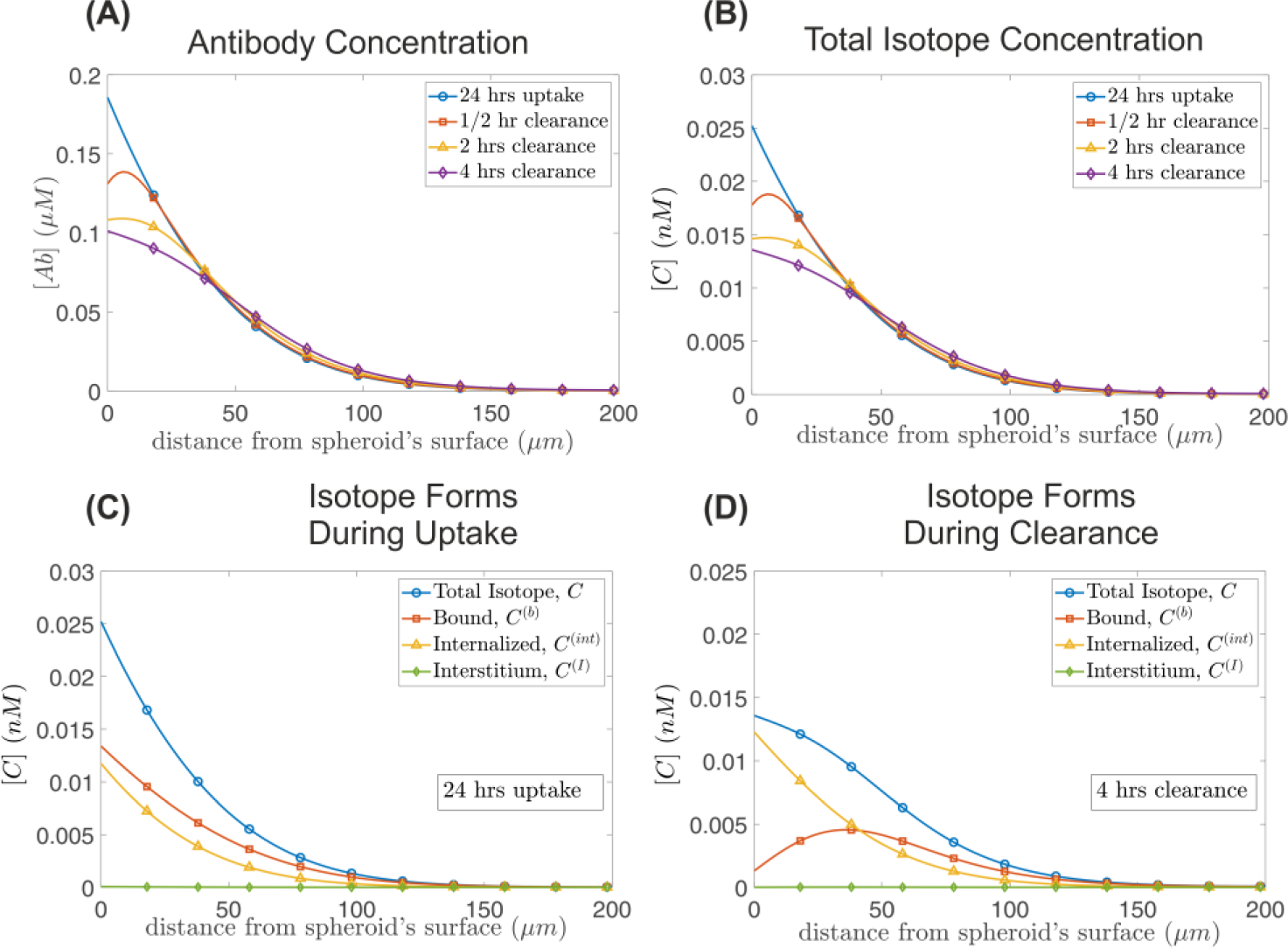
Antibodies isotope carrier simulation. **(A)** Spatial distribution of antibody/isotope complexes within a tumor spheroid at the end of incubation (24 hrs: blue line with open circles), 1/2 hr after incubation (red line with open squares), 2 hrs after incubation (yellow line with open triangles), and 4 hrs after incubation (purple line with open diamonds). **(B)** Isotope concentration spatial distribution at the end of incubation (blue line with open circles), and 1/2, 2, 4 hrs after the completion of incubation depicted with red line and open squares, yellow line and open triangles, and purple line with open diamonds, respectively. **(C)** Spatial distribution of isotope concentration after the completion of the 24 hrs incubation experiment. The total isotope concentration is depicted with a blue line with open circles; receptor-bound isotope concentration is shown in red line and open squares. The internalized isotope concentration is depicted with yellow line and open triangles; and the isotope found in the interstitium is illustrated with the green line with open diamonds. The corresponding profiles of the various forms of isotope during clearance, and specifically 4 hrs after the completion of incubation, are depicted in **(D)**. Parameter values for the simulation: *D_Ab_* = 6 × 10^−12^ *m*^2^ · *s*^−1^, *K_D_* = 5 nM, *k_off_* = 4 × 10^−3^ *s*^−1^, *k_int_* = 1.4 × 10^−5^ *s*^−1^, *P_Ab,up_* = 2.5 × 10^−10^ *m* · *s*^−1^, *P_Ab,cl_* = 8 × 10^−7^ *m* · *s*^−1^, and *R_T_* = 1060 nM. The simulations are performed considering a fixed concentration of antibodies in the solution where antibodies are immersed during incubation: [*Ab*^(*sol*)^] = 240 nM.

### Modelling cocktail isotope delivery, jointly using liposomes as well as antibodies

As reported above, isotope delivery with tumor responsive, isotope releasing liposomes can penetrate and deliver higher values of isotope concentrations further into the spheroid’s interior, whereas antibodies primarily target the exterior regions of a spheroid. It is thus reasonable to expect that the combination (cocktail) of the two carriers results in more uniform isotope distribution within spheroids. Instead of incubating spheroids in a solution with initial radioactivity concentration of 3.7 MBq/L (or equivalently ∼0.0019 nM) and transport the isotope with a single carrier (either liposomes only or antibodies only), we split the isotope to 0.00095 nM encapsulated in liposomes, and 0.00095 nM contained in antibodies. We simulate spheroid incubation with a protocol starting with antibodies for 24 hrs, and liposome incubation for the last 6 hrs of the 24. This incubation schedule is chosen to (approximately) match our knowledge of the corresponding carrier blood clearance half-lives in mice^10^; it is schematically illustrated in the right panel of Fig. 4(A). In the left and center panels of Fig. 4(A), we present the isotope distribution when administered *in equal amounts* utilizing both liposome and antibody carriers. The cocktail of carriers combines the increased isotope penetration capability of liposomes, and the increased concentration level of isotope that antibodies provide at the exterior regions of the spheroids. The model is augmented by a quantification of cell killing described in the “Killing Efficacy” part of the Methods section.

**Fig. 4:**
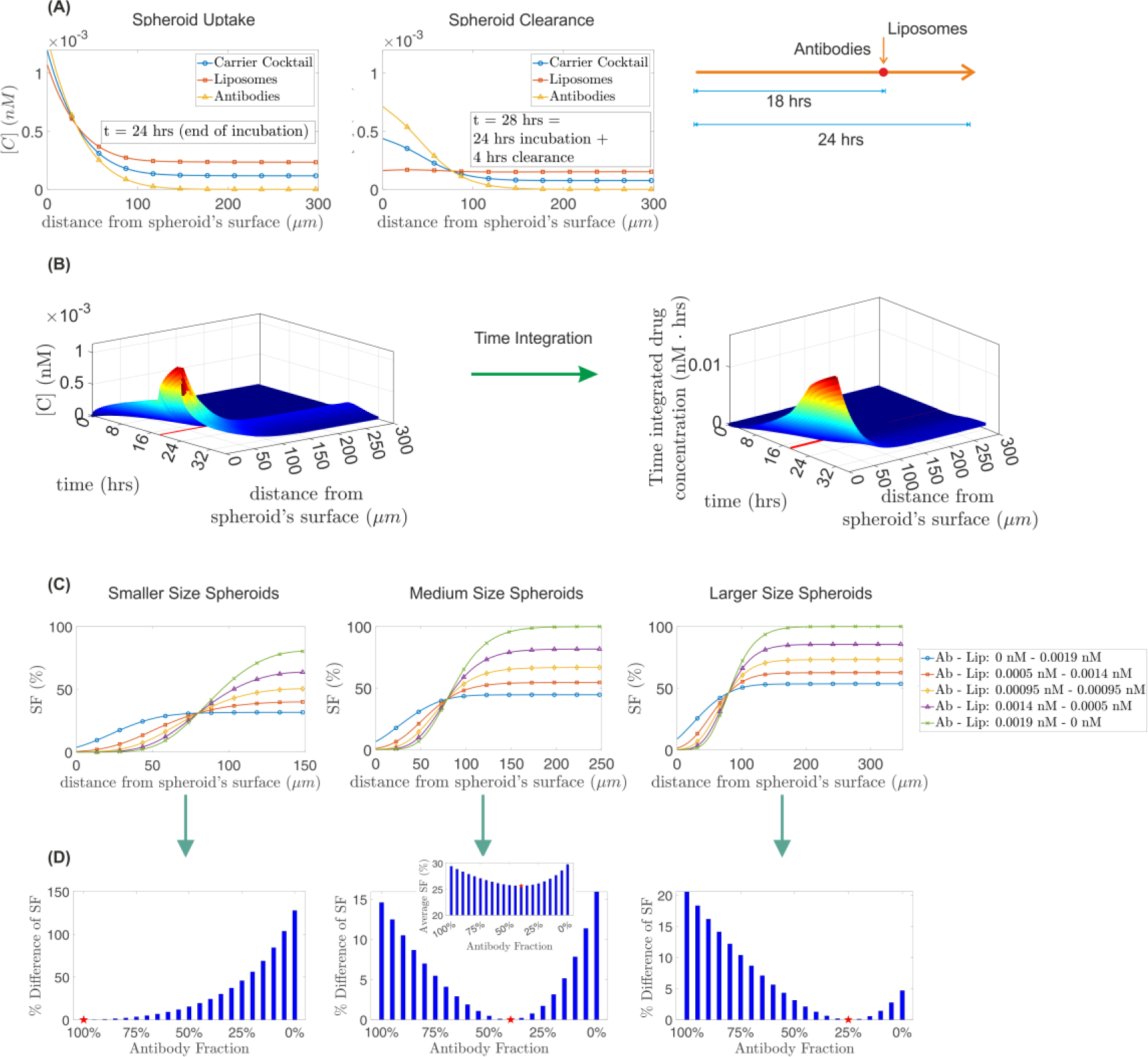
Quantification of treatment efficacy for different isotope carrier combinations. (A) Comparison of isotope concentration spatial profiles obtained for different carrier cocktails. Left and middle panels illustrate profiles at the end of incubation and after 4 hrs of clearance time, respectively. The blue line with circles corresponds to isotope administration with a cocktail of antibody *and* liposome carriers; the red line with squares, and the orange line with triangles represent isotope profiles when using exclusively liposomes and antibodies, respectively. In all cases, the concentration of isotope in the solution where spheroids are immersed during uptake is 0.0019 nM. In the cocktail simulation, liposomes’ and antibodies’ cargo has a concentration of 0.00095 nM each, in the media. The right panel illustrates the isotope delivery temporal policy, with spheroids incubated in a solution of antibody isotope carriers for 24 hrs and liposomes injected in the solution at t=18 hrs (total incubation time with liposome carriers= 6 hrs). **(B)** Left panel: Spatiotemporal evolution of ^225^Ac when carried with a 50/50 % combination of antibodies and liposome carriers. Spheroids are incubated in a solution of antibodies carrying 9.5 × 10^−4^ nM of isotope for 24 hrs. At t=18 hrs liposomes carrying 9.5 × 10^−4^ nM of isotope are administered in the solution. The right panel depicts the time integrated isotope concentration. **(C)** Spatial distribution of the cancer cell *SF* at t=32 hrs for different carrier combinations and different spheroid sizes. Smaller, medium, and larger spheroids have radii of 150 μm (left), 250 μm (center) and 350 μm (right), respectively. *SF* is computed using equation (12) and setting *k_kill_* = 400 *nm*^−1^ · *hrs*^−1^. **(D)** The overall efficiency of different carrier combinations is quantified by illustrating the percentage difference in average *SF* relative to the most effective carrier scheme, indicated by a red star in each tumor size scenario. Higher efficiency for smaller size spheroids is attained when using exclusively antibodies (*SF*=8.1%). For medium size spheroids, optimal efficiency is achieved with a 40% Antibodies/60% Liposomes carrier combination (*SF*=25.7%). The inset shows the *SF* variation for different antibodies/liposomes carrier combination. For spheroids of 350 μm radius, the minimum *SF*=36.8% is computed when using 75% Liposomes / 25 % antibodies.

**Fig. 5:**
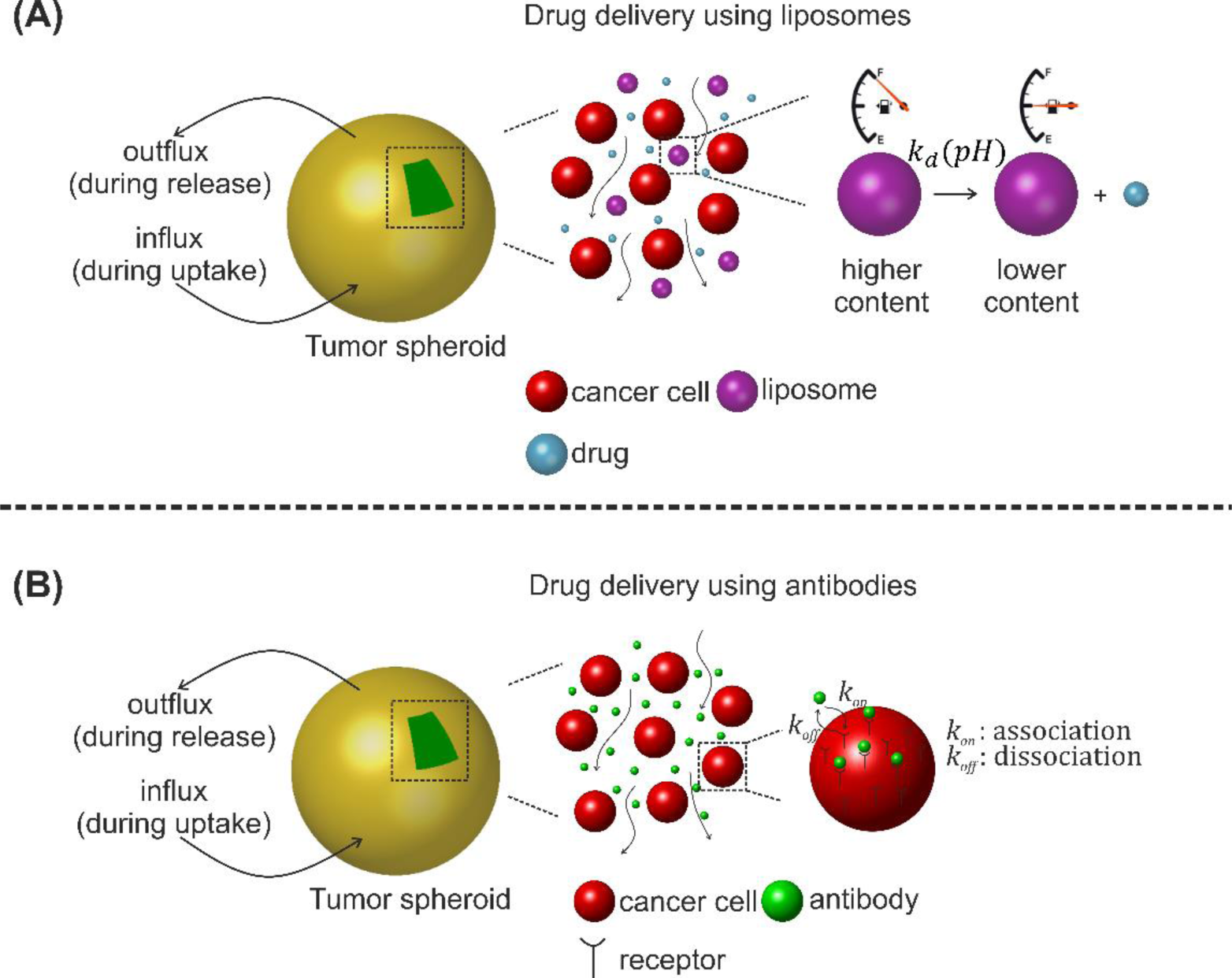
Schematic representation of processes involved during uptake and clearance experiments of isotope-carriers to a spheroid. (A) When the isotope-carriers are liposomes, they infiltrate the spheroid, and release their isotope content with a *pH* dependent rate constant, *k_d_*(*pH*). **(B)** When antibodies are used as isotope-carriers, they diffuse through the interstitium of the tumor and associate/dissociate on/from the cancer cell surface at rate constants *k_on_*, *k_off_*, respectively.

By computing the isotope spatiotemporal distribution (Fig. 4(B) left panel), we can quantify the isotope’s delivered (and remaining) radioactivity explicitly via the time-integral of isotope concentration (Fig. 4(B) right panel) using equation (12). We refer the reader to subsection Killing Efficacy in which we quantify the killing action of the isotope (^225^Ac). In Fig. 4(C) we present the survival fraction (*SF*) of cancer cells’ spatial distribution at time *t* = 32 hrs applying equation (12) and by setting, *k_kill_* = 400 *nm*^−1^ · *hrs*^−1^ = 5.7 × 10^−11^ 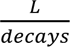. Observe that the overall treatment efficiency when the radius of spheroids is 150 μm (small spheroids) is in general higher, compared to larger size spheroids. In addition, observe the significantly higher efficiency of antibody mediated treatment in the outer regions of spheroids, which drops in the inner regions of spheroids due to the low penetration ability of antibodies. Especially in medium and large size spheroids, there exists a significant portion of the spheroid inner region where the cancer cells remain intact when the isotope is exclusively carried by antibodies. On the other hand, liposome treatment mediates the transport of isotope towards interior regions of the spheroids. Increasing the isotope dose delivered by liposomes makes treatment more effective for larger spheroids; in the absence of antibody carriers; however, the therapeutic efficiency at the outer parts of the spheroids is limited. Clearly, one expects an optimum to develop.

The combined action of antibody *and* liposome carriers is expected to produce higher therapeutic efficacy, and in fact we quantify this through computing the average survival fraction of cancer cells. Figure 4(D) shows the relative difference of (the average) survival fraction with respect to the scheme exhibiting the best performance for small, medium, and large spheroids (left, middle, and right panel, respectively) and different combinations of antibody and liposome carriers. Antibody fraction 100% corresponds to isotope delivery exclusively using antibodies (0.0019 *nm* delivered with antibodies). A fraction 50% denotes a scheme with 0.00095 nM of ^225^Ac being delivered by antibodies and 0.00095 nM is delivered using liposomes. Finally, 0 % corresponds to isotope delivery using exclusively liposomes (0.0019 nM delivered with liposomes). In all cases, the total ^225^Ac isotope concentration in the solution at the beginning of incubation is 0.0019 nM. *Optimal efficiency* is achieved for 100 % antibody fraction for small spheroids (150 μm radius), 40 % for medium size spheroids (7.6 × 10^−4^ nM delivered with antibodies, 11.4 × 10^−4^ nM isotope delivered with liposomes), and 25 % antibody fraction in larger size spheroids (4.75 × 10^−4^ nM delivered with antibodies, and 14.25 × 10^−4^ nM isotope delivered with liposomes). Clearly the model is successful in reproducing, quantifying, and mechanistically validating the initial *in vivo* observations and the intuition behind the success of transport cocktails.

## Discussion

The presented work addresses a critical challenge in the treatment of advanced solid tumors: the delivery heterogeneity of therapeutic agents within the tumor microenvironment. Beyond the intrinsic cellular heterogeneity of tumors, the delivery of therapeutic drugs to different tumor regions varies, leading to nonuniform drug concentrations and doses. By employing two distinct carriers, tumor-responsive liposomes and antibody-drug conjugates, each with complementary delivery properties, leads to improved drug spreading within solid tumors^5,7,10^. While the liposomes release their payload in the interstitium, penetrating deeper tumor regions, the antibodies target perivascular areas. Here, we integrate first-principles reaction-transport models with experimental data to elucidate the synergistic behavior of these carriers. The mathematical models, informed by experiments, successfully reproduce the geographically complementary behaviors observed in the delivery of alpha-particles using carrier cocktails.

The study also explores the optimal synergy between the two carriers for different tumor sizes, emphasizing the importance of tailoring (optimizing) delivery protocols. The quantification of treatment efficacy for various carrier combinations, including exclusive use of liposomes or antibodies, highlights the superior performance of the carrier cocktail approach. The results suggest that an optimum combination of liposomes and antibodies can significantly enhance therapeutic efficacy, offering a potential solution to the delivery heterogeneity challenge in treating established solid tumors. Overall, this work provides valuable insights into the design and optimization of carrier cocktails for effective and uniform drug delivery in the complex landscape of solid tumors.

## Methods

Our reaction-diffusion-type transport model consists of a set of coupled partial differential equations which describe the spatiotemporal evolution of the concentrations of various forms of each moiety (antibodies or liposomes) and the isotope they carry.

### Isotope delivery using liposomes

When a spheroid (our surrogate of tumor avascular regions) is immersed in a solution with constant liposome concentration, then Liposomes (L) are transported through diffusion in the spheroid’s interstitium, and release their isotope-content, *C*, which in turn diffuses in the spheroid interstitium.

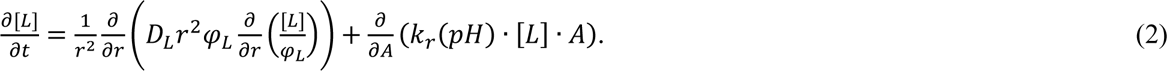

Here, [*L*] = [*L*](*A*, *r*, *t*) denotes the liposome concentration in the interstitium, and depends on the isotope content, *A*, on the radial distance from the tumor spheroid centre, *r* and of course on time, *t*; *k_r_* = *k_r_*(*pH*) denotes the isotope release rate from the liposomes, and is a function of local *pH*. *D_L_* is the effective diffusion coefficient of liposome in the spheroid, and *φ_L_* denotes the fraction of spheroid volume accessible to liposomes.

At the spheroid’s external surface, the mass flux rate is prescribed from the following *mass-transfer relation*:

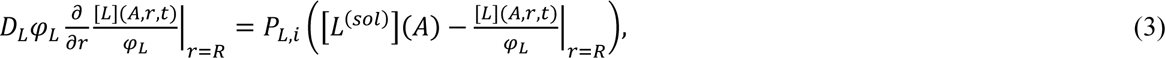

where *P_L,i_* is the mass transfer coefficient for liposomes during uptake (when *i* = *up*) and during clearance when (*i* = *cl*; [*L*^(*sol*)^])(*A*) is the concentration of liposomes with isotope content *A* in the solution in which spheroids are immersed. *R* denotes the radius of the spheroid. In the solution liposomes with isotope content *A*=100% are initially present.

Isotope release and transport in the spheroid is governed by:

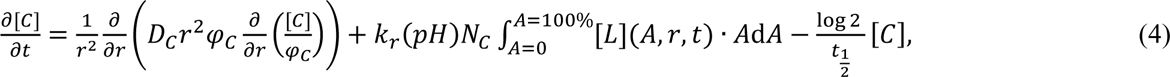

where [*C*] is the isotope concentration in the interstitium, *D_C_* is the effective diffusion coefficient of the isotope in the tumor interstitium, and *φ_c_* denotes the fraction of spheroid accessible to isotope. *N_C_* is the ratio of isotope content per liposome (isotope mol/liposome mol) when *A* = 100%. In all simulations presented, this maximal isotope content per liposome is *N_C_* = 0.017 moles isotope / moles liposome. Finally, 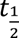 denotes the half-life of the radioactive isotope (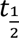 =10 days for ^225^Ac). At the exterior surface of the spheroid, *r* = *R*, we prescribe the flux of isotope by continuity:

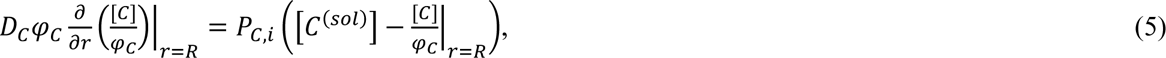

where *P_C,i_* denotes the mass transfer coefficient for isotope from the solution to the spheroid during uptake, *i* ≡ *up*, and during clearance, *i* ≡ *cl*, respectively. [*C*^(*sol*)^] is the concentration of the free isotope in the solution where the spheroids are immersed. Since the solution initially contains only isotope encapsulated by liposomes, [*C*^(*sol*)^](*t* = 0) = 0, and we assume here that, for a large bath, it remains practically negligible during incubation.

### Isotope delivery using specific antibodies

When a spheroid is immersed in a solution with constant antibody concentration, antibodies enter the spheroid through the outer tumor surface. Once in the tumor interstitium, antibodies are transported by diffusion, bind with surface receptors, dissociate from them, and/or become internalized within the cells. Again, we formulate reaction-diffusion equations that describe the processes involved, and in particular:

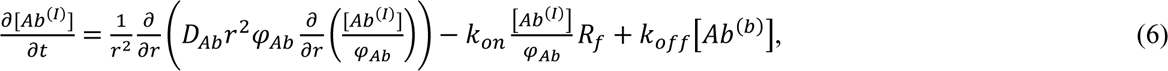

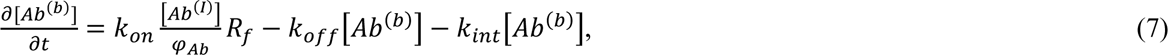

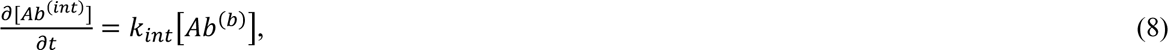

where [*Ab*^(*I*)^] denotes the antibody concentration in the interstitium, [*Ab*^(*b*)^] is the concentration of the antibody/receptor complex, and [*Ab*^(*int*)^] denotes the concentration of internalized antibody; *D_Ab_* denotes the effective diffusion coefficient of antibodies when transported in the interstitium; *k_on_*, *k_off_* denote the association and dissociation rate constants on/from the cell surface, respectively and *k_int_* is the internalization rate constant; *R_f_* is the concentration of unoccupied (by antibodies) receptors, and *φ_Ab_* is the fraction of spheroid volume accessible to antibody.

The mass flux rate at the spheroid’s external surface is given by:

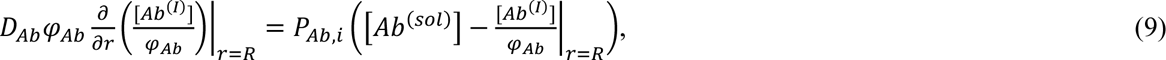

where [*Ab*^(*sol*)^] is the antibody concentration in the solution in which spheroids are immersed, and *P*_*Ab*,*i*_ denotes the mass transfer coefficient, equal to *P_Ab,up_* for uptake experiments, and *P_Ab,cl_* for clearance experiments. Finally, we consider the following material balance for surface receptors:

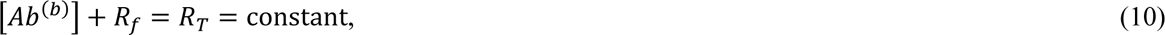

with *R_T_* denoting the initial (unbound) receptor concentration. All concentrations are expressed over the total spheroid volume (both accessible and inaccessible to antibodies).

### Killing efficacy

The cargo-isotope is an *a* −particle generator which can result in high cancer-cell killing with minimal irradiation. Here, we consider ^225^Ac which decays with a 9.9-day half-life, generating a total of four *a* −particles, with range in tissue between 40 and 80 µm (5-10 cell diameters), and three radioactive daughters^13^. Without loss of generality, the decay of radioactive daughters is assumed to occur in the vicinity of the parent ^225^Ac nucleus. In this study, we assume that the death kinetics of the cancer cells in the spheroid follow:

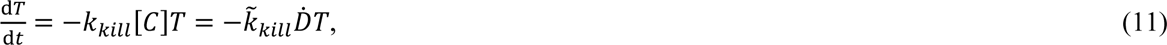

where *T*(*r*, *t*) denotes the density of surviving cancer cells at radial distance, _r_, from the center of and time, *t*, [*C*] is the local concentration of ^225^Ac, and *k_kill_* denotes the killing rate constant of cancer cells (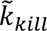 when death kinetics are expressed in terms of decay rate, 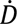). The concentration of ^225^Ac isotope can be correlated with its decay rate, given the half-life of the isotope, 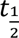 = 9.9 days, and that each decay generates four alpha particles and three radio-active daughters^13^. The radioactivity of ^225^*Ac* is ∼2,000 *MBq*/*nmol*, thus 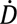; ≅ 2,000 × [*C*] MBq/L (when [*C*] is expressed in nM). Then, a formula providing the survival fraction (*SF*) of cancer cells follows:

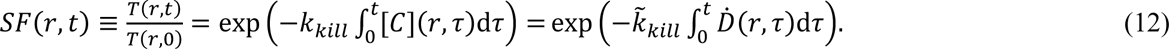

In conclusion, the killing efficiency of different isotope carrier combinations is measured by the survival fraction of cancer cells at each radial distance, *r*, and each time instance, *t*.

### Experimentally informed parameter fitting

#### Porosity Profiles

Multicellular, 3D spheroids of BT474 breast cancer cells, overexpressing the HER2-targeted marker, were formed using established methods^4^, and were utilized as the surrogates of solid tumors’ avascular regions. The interstitial pH gradient in these spheroids was previously measured, using a fluorescent pH indicator, and was shown to range from approximately 6.5 in the core, to 7.4 at the edges, when suspended in regular media (see also Fig.(S-1) of the Supplementary Material). To quantify the spatiotemporal profiles of each modality, spheroids were removed from the incubation suspension at different time points (as described in detail in references^5,7,10,12^), were snap frozen, sliced, and the equatorial sections were imaged by a fluorescence microscope. The average radial intensities of each fluorescently-labelled modality were calculated using an in-house erosion MATLAB code, and the radial distributions were quantified by comparing, for each fluorescent species, to calibrated curves, generated using the same microscope in a cuvette, of pathlength identical to the thickness of the spheroid sections, containing known concentrations of each fluorescent species/moiety^5,7,10,12^. For the derivation of each moiety’s porosity profiles, we worked with the non-reacting/binding corresponding moieties: a non-specific antibody, Rituximab (that was shown to not specially bind to BT-474 cells (see SI Figure S1), and with non-releasing, non-adhering liposomes (as characterized in^7–9^). Data were collected by incubating spheroids for long times (longer than 24 hours) until the concentration profiles of liposomes and antibodies remained practically unaltered. In both cases, the steady-state solution of the model (see equations (2) & (6)) must be a uniform solution, and in particular: [*L*]^(*steady*)^(*r*) = *φ_L_*[*L*^(*sol*)^] and [*Ab*]^(*steady*)^(*r*) = *φ_Ab_*[*Ab*^(*sol*)^] for liposomes and antibodies, respectively. Thus, one could infer the porosity profile by fitting curves *φ_L_*(*r*), *φ_Ab_*(*r*) to the data, [*L*]/[*L*^(*sol*)^] and [*Ab*^(*I*)^]/[*Ab*^(*sol*)^] (see Figs 1(A) & (B)). For the computation of porosity profile, spheroids were incubated with ’’drug/isotope-empty’’, non-adhering liposomes, and non-specific, non-associating antibodies Rituximab (*k_on_* = *k_off_* = 0, see equation (6) in the Methods section). The resulting porosity fitting for non-adhering, non-releasing liposomes is: *φ_L_* ≈ 0.44*r*^3.2^ + 0.56, and for non-specific antibodies: *φ_Ab_* ≈ 0.83*r*^5.21^ + 0.17. The porosity profile for liposomes features larger variation (compared to antibodies), with considerably lower values at the spheroid’s centre; at the centre of a spheroid only ∼17 % of the volume is accessible to liposomes.

#### Obtaining transport parameters for non-adhering, non-releasing liposomes

Estimation of transport parameters, *D_L_*, and *P*_*L,up*(*cl*)_ is performed by fitting the continuum model (equations (2) & (3)) to experimental measurements of liposomal radial distributions during uptake and clearance experiments. We simulate the spatiotemporal evolution of *non-adhering, non-releasing, isotope-free* liposomal carriers, which simplifies our computations. The experimental data are obtained by incubating spheroids in a solution of concentration, [*L*^(*sol*)^] = 0.5 mM liposomes for 6 hrs; then spheroids are fished from the medium and are immersed in clear media (clearance) for another 24 hrs. Figure 6(C) illustrates the best fitting simulation of the continuum model, equations (2) & (3) on experimental measurements of non-adhering, non-releasing liposomes. Best fits are obtained using MATLAB’s function *nlnfit*, which performs nonlinear regression using iterative least squares estimation. The estimated effective diffusion coefficient of liposomes in the spheroids is: *D_L_* = (1.46 ± 0.13) × 10^−13^ *m*^2^ · *s*^−1^. The mass transfer coefficient of liposomes during incubation is estimated as: *P_L,up_* = (1.91 ± 0.18) × 10^−9^ *m* · *s*^−1^. The mass transfer coefficient of liposomes during clearance experiments (spheroids immersed in clean water) is estimated: *P_L,cl_* = (5.81 ± 1.0) × 10^−9^ *m* · *s*^−1^.

**Fig. 6:**
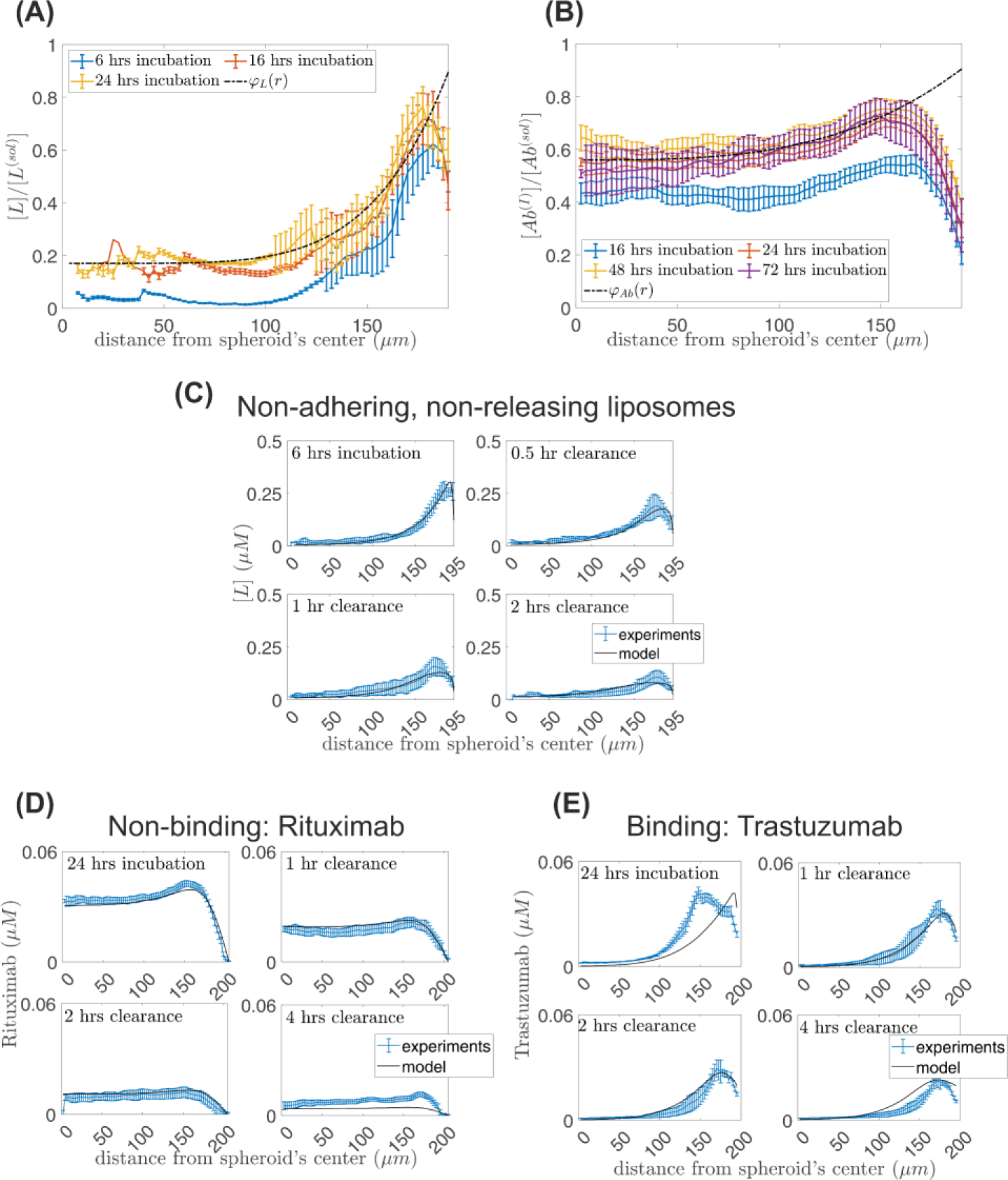
**Fitting of transport and kinetic properties**. **(A)** Porosity profile for non-adhering liposomes. **(B)** Porosity profile for non-binding antibodies. **(C)** Fitting of transport properties on non-releasing, non-adhering liposomes uptake and clearance experiments. **(D)** Fitting of transport properties on non-binding, Rituximab antibody experiments (uptake and clearance). **(E)** Fitting of kinetic properties on binding/specific Trastuzumab antibodies experiments.

Similarly, to estimate the diffusion coefficient and mass transfer coefficients of the drug released from liposomes within the interstitium, spheroids were incubated with 3 µM free Newport Green (NG) for up to 6 hours and were sampled both during the uptake and clearance of NG, at different time points. The spatiotemporal distributions of NG within spheroids were quantified and were analysed as described above for liposomes. The diffusion coefficient for NG is estimated: *D_C_* = (1.96 ± 0.19) × 10^−11^ *m*^2^ · *s*^−1^, the drug mass transfer coefficient during uptake and clearance: *P_C,up_* = (1.7 ± 0.01) × 10^−8^ *m* · *s*^−1^ and *P_C,cl_* = (2.9 ± 0.2) × 10^−8^ *m* · *s*^−1^, respectively. Assuming that the mass transport property values of NG are similar to those of the isotope, we incorporate them into our computations.

#### Transport and kinetic parameters for Antibodies

We first estimate transport properties of antibodies by fitting equations (6) - (9) to experimental data of non-binding antibodies (Rituximab). Tumor spheroids are immersed in a 0.06 μM Rituximab solution for 24 hrs (incubation/uptake experiment); then the spheroids are fished from the medium and are immersed in clear media (clearance) for another 24 hrs. The best fitting is illustrated in Fig. 6(D) for *DAb* = (8.38 ± 0.41) × 10^−12^ *m*^2^ · *s*^−1^. The uptake mass transfer coefficient is estimated as: *PAb*,*up* = (1.54 ± 0.04) × 10^−9^ *m* · *s*^−1^, and the release mass transfer coefficient of antibodies is computed: *PAb*, *cl* = (8.58 ± 0.31) × 10^−9^ *m* · *s*^−1^.

To estimate the association, dissociation and internalization rate constants (*k_on_*, *k_off_*, *kin*, see equation (6)), we fit the continuum-level model, equations (6)-(9) on experimental measurements of the binding/specific antibody, the HER2-targeting Trastuzumab. Trastuzumab diffuses in the spheroid through the interstitium, and also associates (dissociates) on (from) the surface of cancer cells and internalizes; here, the porosity, *φ_Ab_*, diffusion coefficient value, *D_Ab_* and mass transfer coefficients, *P*_*Ab,up*(*cl*)_, are adopted from the non-binding antibody (Rituximab) experiments. The total concentration of antibody receptors, *R_T_* is estimated from experimental measurements, and *R_T_* ≈ 1060 nM. The best fitting simulations of the Trastuzumab uptake and release experiments are illustrated in Fig. 6(E) for *k_off_* = (4 ± 1.6) × 10^−3^ *s*^−1^, 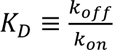 = 6.76 ± 1.75 nM (the equilibrium dissociation constant between the antibody and its antigen), and *kint* = (1.44 ± 0.57) × 10^−5^ *s*^−1^ (the internalization rate constant of the antibody). Our computed values for antibodies transport and kinetic parameters compare reasonably well with reported values in the relevant literature (see Table 1).

**Table 1.** List of kinetic and reaction parameters for Rituximab and Trastuzumab antibodies.

| Parameter | Fitting | Typical Values |
| --- | --- | --- |
| $D_{Ab} (\mu m^2 \cdot s^{-1})$ | $8.38 \pm 0.41$ | $5-50^{18-21,19,20,21}$ |
| $P_{Ab,up(cl)} (m \cdot s^{-1})$ | $2.5 \times 10^{-10} - 8 \times 10^{-7}$ | $3 \times 10^{-9} - 1.5 \times 10^{-7} \text{ }^{22,23}$ |
| $K_D$ (nM) | $6.76 \pm 1.75$ | 1-10 (often in IgGs)<sup>24</sup> |
| $k_{off} (s^{-1})$ | $(4 \pm 1.6) \times 10^{-3}$ | $10^{-6} - 4 \times 10^{-3} \text{ }^{24,25}$ |
| $k_{int} (s^{-1})$ | $(1.4 \pm 0.57) \times 10^{-5}$ | $2 \times 10^{-6} - 2 \times 10^{-4} \text{ }^{26,27,28,29}$ |

## Data Availability

The data that support the findings of this study are available within this article and its Supplementary Information. Further data are available from the corresponding authors upon reasonable request.

## Code Availability

The codes used for this work can be found in: https://github.com/mihaliskavousanakis/TransportCocktails.

## Acknowledgements (optional)

S.S. acknowledges the financial support of W.W. Smith Charitable Trust, the Allegheny Health Network-Johns Hopkins Cancer Research Fund, Maryland Innovation Initiative – TEDCO. Y.K. was supported from the US Air Force Office of Scientific Research.

## Supplementary Information

Supplementary Information: A. Interstitial pH profile in spheroids, B. Dependence of liposomal drug release on pH, C. Binding parameters for specific antibody with BT-474 cells.

