## Supplementary Information for "Transport Cocktails for Cancer Therapeutics"

### Supplementary Material


#### Michail E. Kavousanakis, Omkar Bhatavdekar, Remco Bastiaannet, Yannis Kevrekidis, Stavroula Sofou

Affiliations

**School of Chemical Engineering, National Technical University of Athens, Iroon Polytechneiou 9, Athens,15780, Greece**

Author-1

**Department of Chemical and Biomolecular Engineering, Institute for NanoBioTechnology, Johns Hopkins University, Baltimore, Maryland 21218, USA**

Author-2, Author-4 & Author-5

**Department of Radiology and Radiological Science, School of Medicine, Johns Hopkins University, Baltimore, Maryland 21287, USA**

Author-3


#### A. Interstitial pH profile in spheroids

Multicellular spheroids serve as tumor surrogates, specifically emulating the avascular region within the tumor. These spheroids not only replicate the 3-dimensional architecture and microenvironment found in tumors but also mimic the spatial variation of pH observed in tumors. In Fig. S1, we present experimental measurements of the interstitial pH profile by incubating BT-474 spheroids with the membrane impermeant pH-indicator, SNARF-4F. The results reveal an acidic environment near the interior of the spheroid, while alkaline conditions dominate in the outer zones of the spheroid with a pH $\approx 7.4$.

Incorporating the pH spatial variation into our model involves adopting the following formula:

$$pH = \frac{a-b}{2}\tanh\left(\left(\frac{r}{R} - r_o\right)c\right) + \frac{a+b}{2}. \qquad \text{(S-1)}$$

Here, $a, b, c, r_o$ are fitting parameters obtained through the empirical fitting of Eq.(S-1) to the experimental pH measurements in BT-474 spheroids. The resulting set of parameter values is as follows: $a = 7.49$, $b = 6.58$, $c = 7.87$, and $r_o = 0.624$.

#### B. Dependence of liposomal drug release on pH

The kinetic parameters relevant to liposomes as drug carriers exhibit pH dependency. Utilizing Eq.(S-1), we can describe their spatial variability within spheroids. Specifically, the drug release rate from pH-responsive liposomes follows a linear relationship with pH (refer to Fig.S2(b)). To determine the drug release kinetic constant, $k_d$, we measure the drug's retention percentage, $y$, and fit it to the expression:

$$y = y_\infty + (100 - y_\infty)exp(-k_d t), \quad \text{(S-2)}$$

where $y_\infty$ is the retention percentage under steady-state conditions. Fig. S2(A) illustrates the evolution of drug retention percentage for various pH values covering the range from alkaline to acidic conditions found in spheroids. For each pH value, we fit Eq.(S-2) to compute the drug release constant, $k_r$, and subsequently perform a linear fit of $k_r$with pH as presented in Fig. S2(B). The resulting empirical relationship is expressed as follows: $k_r = 0.48 - 0.064 \cdot pH\ min^{-1}$ .One can observe that drug release is facilitated in acidic conditions, implying an increased drug release rate within the interior regions of the spheroid.

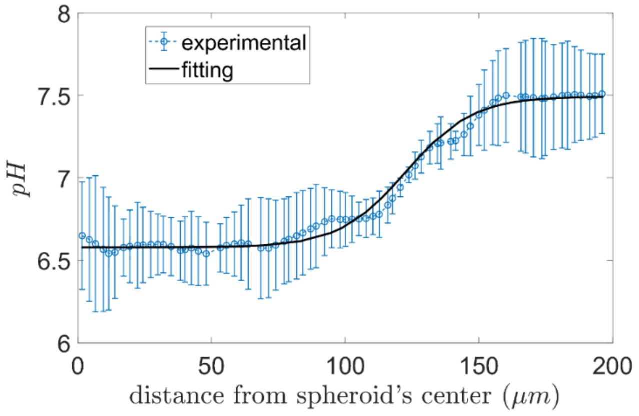


**Fig.S1** Spatial variation of pH in BT-474 spheroids and fitting using Eq.(S-1). Error bars correspond to standard deviation between the means of intensities for n=3-5 spheroids per condition.

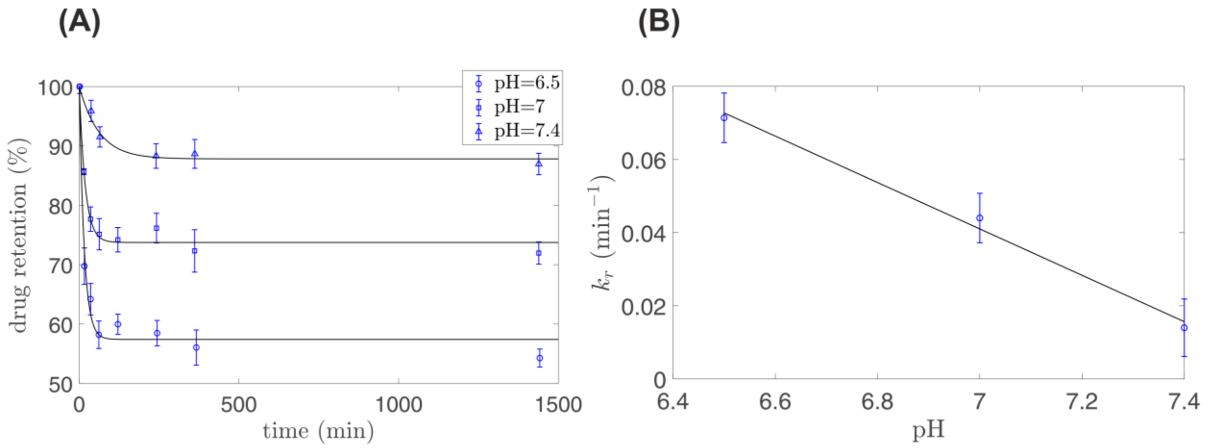


**Fig.S2** Liposome drug release rate kinetics dependence on pH. (A) Evolution of drug's retention in non-adhering liposomes for different pH conditions. For each pH, $k_r$ is computed by fitting Eq. (S-2). (B) Linear fitting of the computed $k_r$ values with pH.

#### C. Binding parameters for specific antibody with BT-474 cells

BT-474 cells were incubated with a range of FITC-labeled antibody (Ab) concentrations in the presence, and absence, of blocked conditions while maintained on ice. After a 90-minute incubation, the cells underwent 3 washes with ice-cold PBS. Subsequently, the cells were resuspended in deionized water, and the fluorescence intensity was measured. Fig. (S-4) displays the measured bound Ab concentration in nM per million cancer cells for different Ab concentration values. The association/dissociation reactions of the specific antibody on cancer cells are:

$$Ab + R_{f,s} \underset{k_{off}}{\overset{k_{on}}{\rightleftharpoons}} Ab^{(b)}. \quad \text{(S-3)}$$

Here, $Ab$ represents the free form of the antibody, $R_{f,s}$ indicates the unbound Ab receptors on the cell surface, and $Ab^{(b)}$ denotes the bound form of the antibody. $k_{on}$ and $k_{off}$ denote the association and dissociation rate constants of the antibody with/from the cell surface, respectively. At equilibrium, the concentration of bound antibody is obtained using the following relation:

$$\left[Ab^{(b)}\right] = \frac{R_{T,s}[Ab^{(sol)}]}{[Ab^{(sol)}]+K_D}, \quad \text{(S-4)}$$

where $[Ab^{(sol)}] = [Ab] + [Ab^{(b)}] \approx [Ab]$ represents the total concentration of Ab in the solution where cancer cells are incubated, and $K_D \equiv k_{off}/k_{on}$ denotes the dissociation constant. $R_{T,s}$ denotes the total Ab receptors on the cell surface, given by: $R_{T,s} = R_{f,s} + [Ab^{(b)}]$. By fitting Eq.(S-4) to the experimental measurements of bound Ab (as seen in Fig. S3), we compute $R_{T,s} \approx 2.12$ nM.

The bound Ab measurements were performed for a suspension containing $10^6$ cancer cells in 1 ml. Therefore, the number of receptors per cancer cell is approximately $1.3 \times 10^6 = \frac{R_{T,s} \cdot N_A}{10^6 \frac{cells}{ml}}$, where $N_A$ is the Avogadro's number. To obtain the concentration of Ab receptors in spheroids, $R_T$, (see Eq. (10)), we need to account for the cancer cell density within the spheroids. For instance, considering BT-474 spheroids with approximately 500,000 cells/mm$^3$, the concentration of Ab receptors is approximately: $R_T \approx 1060\, nM (= 1.3 \times 10^6 \frac{receptors}{cell} \cdot 500{,}000 \frac{cells}{mm^3} \cdot \frac{1}{N_A \frac{receptors}{mole}})$.

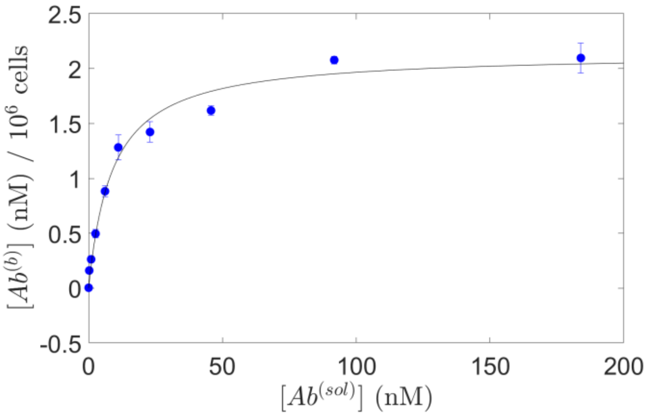


**Fig.S3** Measured bound Ab concentration, $[Ab^{(b)}]$, when BT-474 cells are incubated in a solution with free Ab concentration, $[Ab^{(sol)}]$. The solid black represents the fitting of Eq. (S-4) to the experimental measurements.
